## Supplementary figures and images for "Hundreds of myosin 10s are pushed to the tips of filopodia and could cause traffic jams on actin"

### Fig. 1 supplement 1

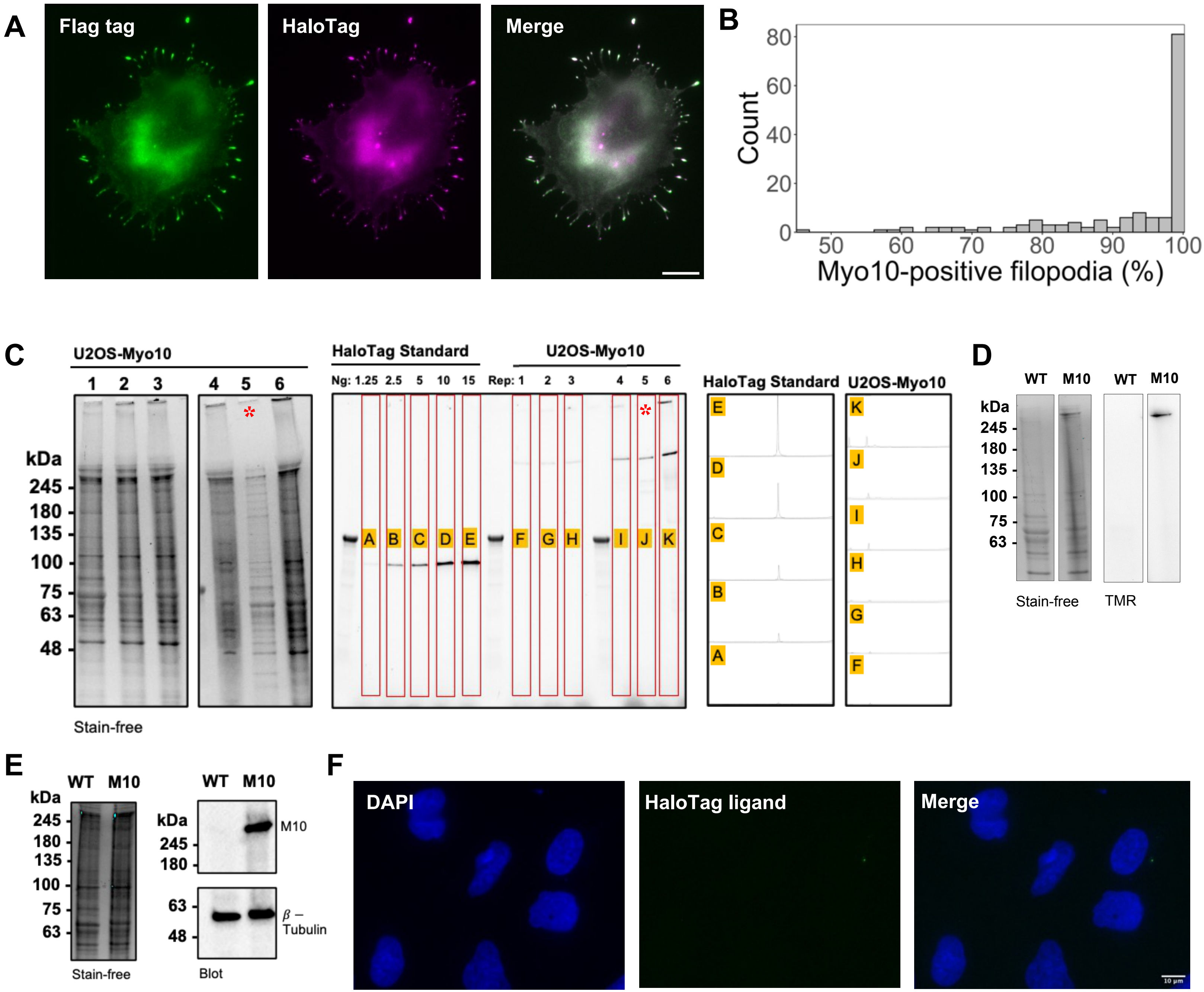

### Fig. 1 supplement 2

A

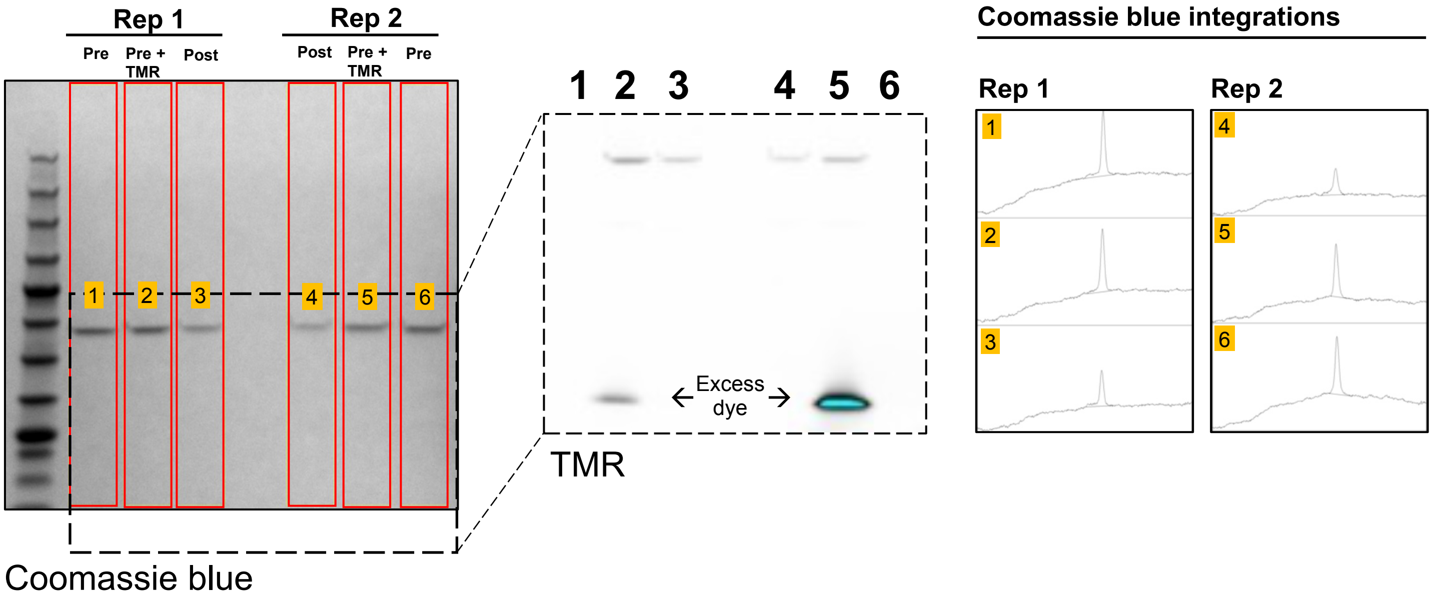


B

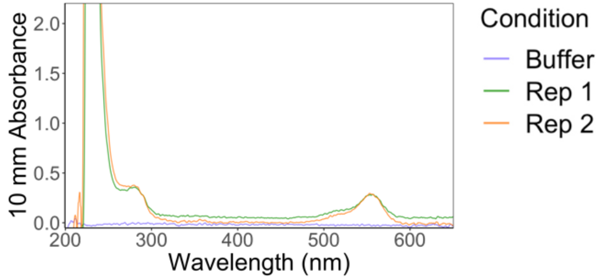


C

| | Rep 1 | Rep 2 |
|---|---|---|
| Pre protein (μM) | 6.8 | 6.8 |
| Pre gel signal | 7925 | 6553 |
| Post/pre gel signal | 0.578 | 0.623 |
| Post Abs553 | 0.2795 | 0.296 |
| Post dye (μM) | 3.6 | 3.8 |
| % dye labeling | 0.912 | 0.8962 |

D

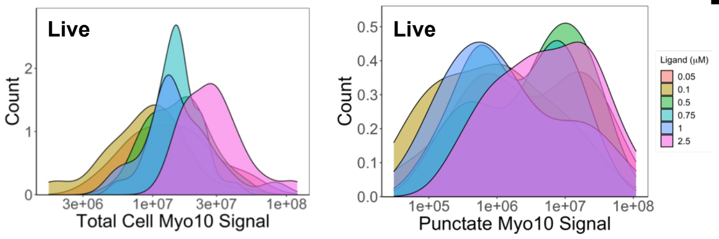


E

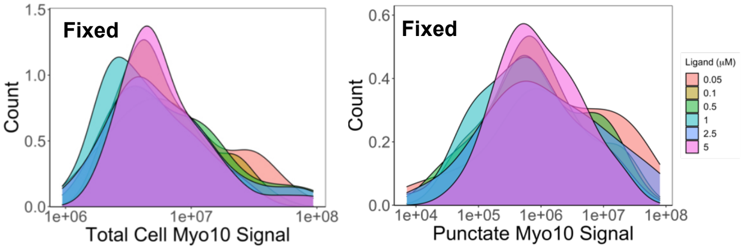

### Fig. 1 supplement 3

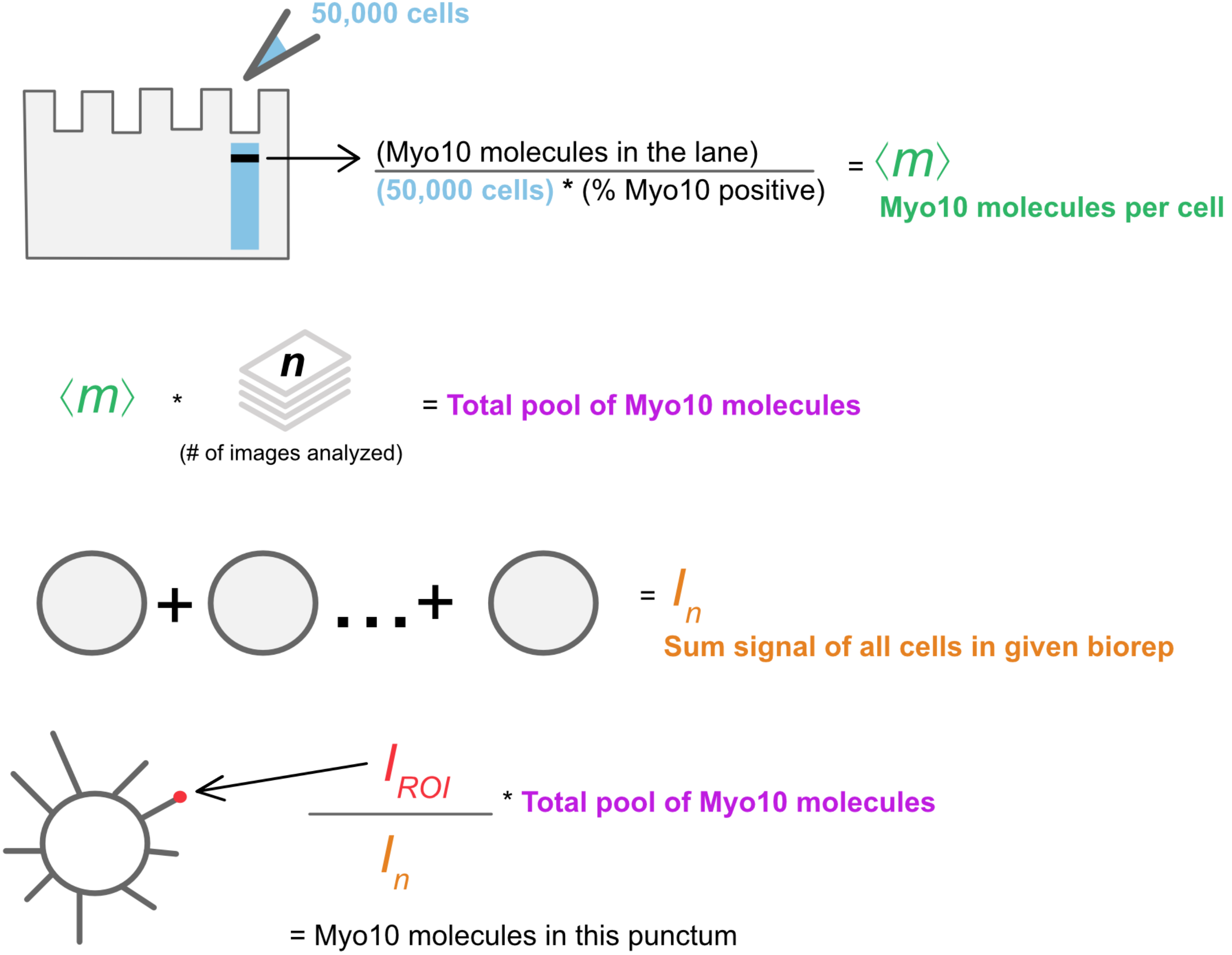

### Fig. 2 supplement 1

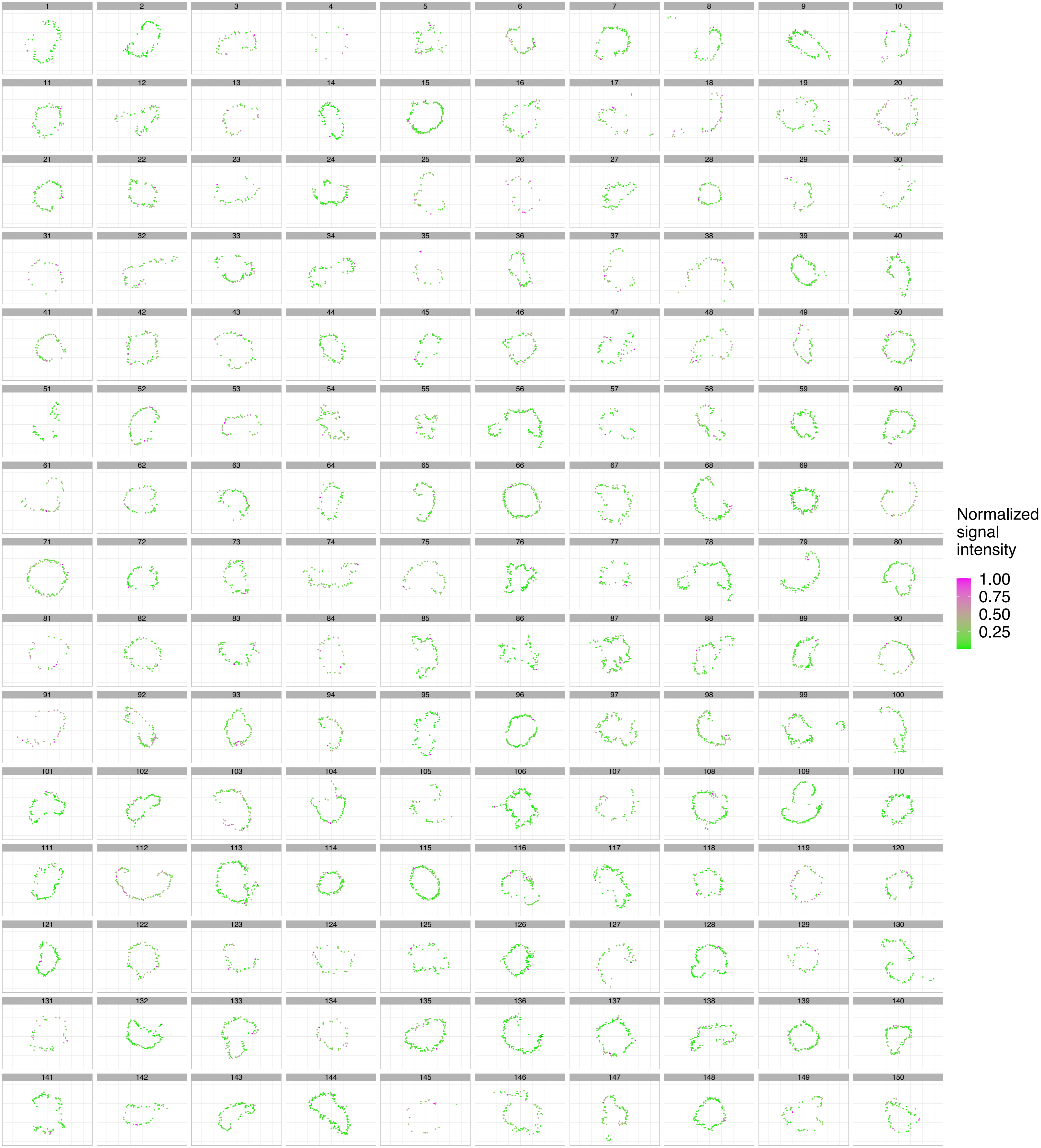

### Fig. 2 supplement 2

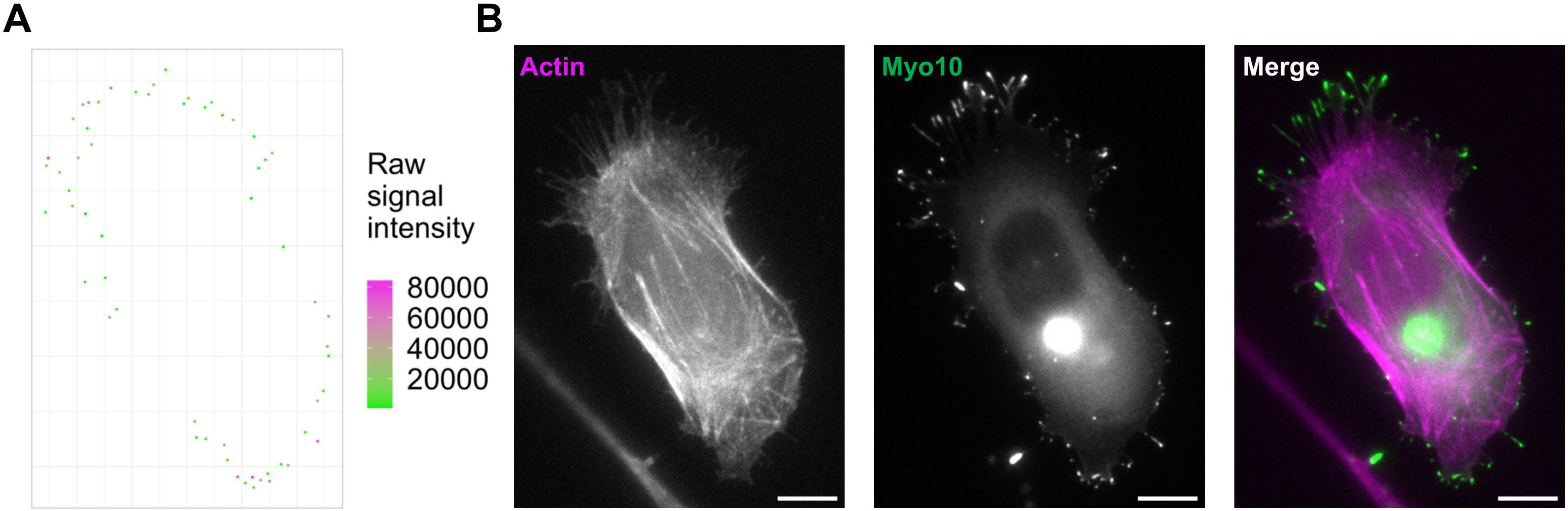

### Fig. 3 supplement 1

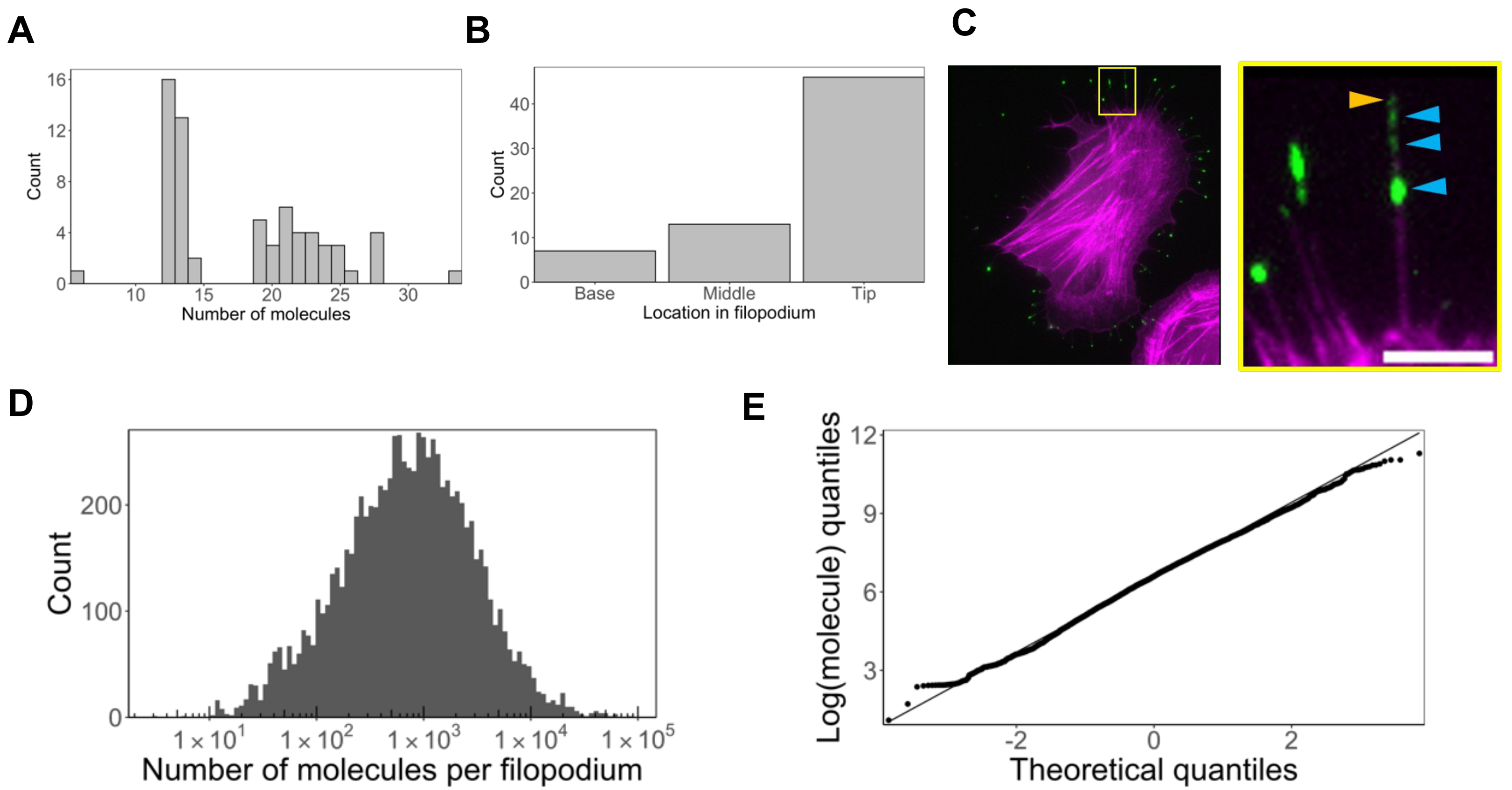

### Fig. 4 supplement 1

A

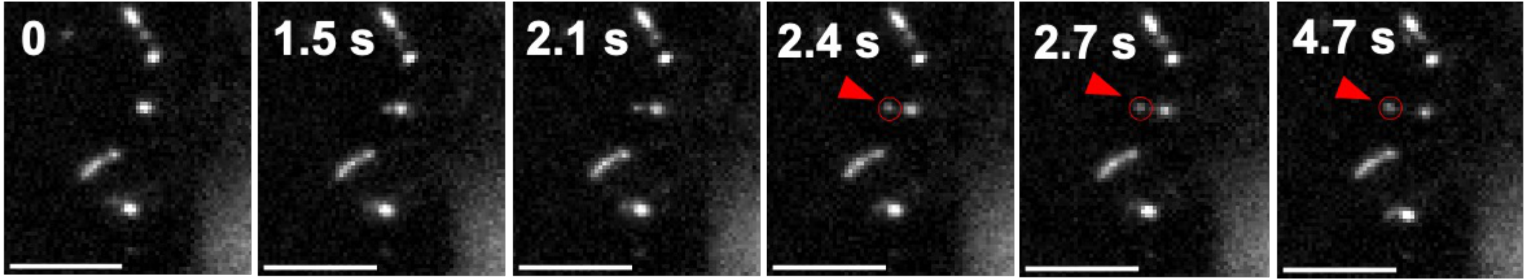
