## Supplementary material for "Hundreds of myosin 10s are pushed to the tips of filopodia and could cause traffic jams on actin": Fig. 3 supplement 2

Aramaki et al. (2016) reported:

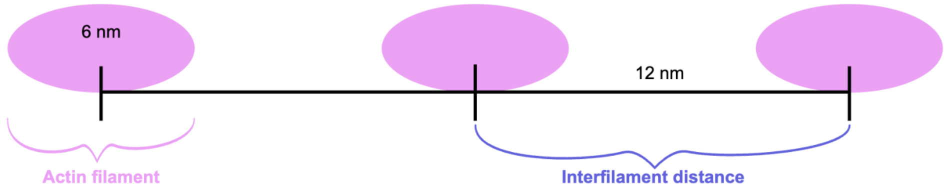


Say a filopodia is ~200 nm in diameter:

$$\frac{200\ \text{nm}}{30\ \text{nm}} \sim 6.7 \text{ actin filaments across a filopodium shaft}$$

Aramaki et al. (2016) found ~30 actin filaments in a filopodium. Because actin filaments are packed into a hexagonal lattice, we can use this model:

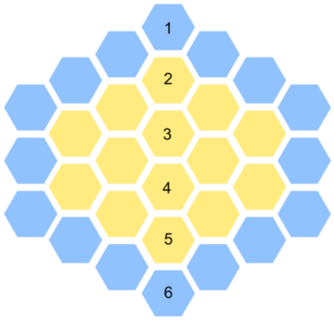


Above, each hexagon represents an actin filament:

- 6 filaments across, 30 filaments in total
- Blue hexagons represent filaments on the outside of an actin bundle in a filopodium
  - Therefore, 16/30 filaments are available for interaction with Myo10

Nagy et al. (2010) posited:

- Knowing that 1 actin filament is composed of 13 monomers per helical turn,
  - Only 4/13 actin monomers are available to Myo10
- Based on that logic:

$$16 \text{ outer actin filaments} * 13 \text{ monomers} = 208 \text{ total actin monomers}$$
$$208 * \frac{4}{13} = 64 \text{ actin monomers available to Myo10 in a helical turn in a filopodium}$$

Assuming 30 actin filaments in a filopodium:

$$30 \text{ total actin filaments} * 13 \text{ monomers} = 390 \text{ total actin monomers in a helical turn in a filopodium}$$

Therefore,

$$\frac{64}{390} = 0.1641$$
$$\sim 16.41\% \text{ of actin monomers are accessible to Myo10 in a filopodium}$$

Zhuralev et al. (2012) modeled the F-actin monomer concentration in a filopodium as:

$$\frac{\text{N}}{\pi * \text{R}^2 * \delta}$$
$$\text{N} = \text{ number of actin filaments}$$
$$\delta = 2.7 \text{ nm; half the size of an actin monomer}$$
$$\text{R} = \text{ radius of filopodium cross-section}$$

If we consider a filopodium containing N = 30 and R = 100 nm:

$$\frac{30 \text{ filaments}}{\pi * 100^2 * 2.7} * \frac{\text{mol}}{6.02 * 10^{23}} * \frac{\text{nm}^3}{10^{-24}} * \frac{10^6\,\mu\text{M}}{\text{M}}$$
$$= 587\ \mu\text{M of F-actin monmers in a filopodium}$$

Based on the earlier calculation that ~16.41% of actin monomers are accesible to Myo10:

$$0.1641 * 587$$
$$= 96\ \mu\text{M of accessible actin monomers to Myo10 in a filopodium}$$
