## Supplementary material for "Hundreds of myosin 10s are pushed to the tips of filopodia and could cause traffic jams on actin": Fig. 3 supplement 3

Consider a filopodial tip, as represented by a hemisphere:

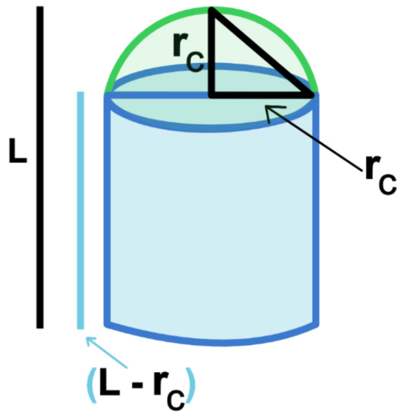


Assume radius of cylinder is 100 nm, which represents the filopodial shaft radius:

$$\text{Surface area of curved hemisphere: } = 2 * \pi * r_c^2 = 62800 \text{ nm}^2$$

In Fig. 3F, we estimate the membrane area available for Myo10 to include the curved hemisphere area AND additional membrane space occupied by the length (L) of the Myo10 spot measured.

$$\text{Surface area of cylinder (area of rectangle): } = 2 * \pi * r_c * (L - r_c) = 2 * \pi * 100 * (L - 100)$$
$$\text{Total membrane surface area considered: } = 200 * \pi * (L - 100) + 62800$$

How many Myo10 molecules can the filopodial tip membrane surface area account for?

- Let's assume Myo10 occupies a rectangular area of the membrane.
- Potentially the PH3 and MyTH-FERM domains of Myo10 contribute to membrane binding. If so, we can use an Alphafold structure prediction of Myo10 PH1, PH2, PH3 and MyTH-FERM domains.
  - Roughly measuring the longest length-wise and width-wise distances in PyMol gives 13.17 nm x 6.99 nm.
  - This provides an estimate of Myo10 area occupancy on the membrane.

$$\text{Estimate of Myo10 membrane area occupancy : } 13.17 * 6.99 = 92 \text{ nm}^2$$

In the case of a hemispherical tip + additional membrane, one can imagine that:

$$[200 * \pi * (L - 100) + 62800]/92 = \text{Myo10 molecules that can bind to hemispherical membrane}$$

This equation is plotted in Fig. 3F.

In reality, the number of Myo10 molecules that can bind the membrane is likely lower, considering that: 1) There would not be perfect packing of the Myo10 molecules on the membrane; and 2) There are other proteins occupying the same space on the membrane.

In some instances, Myo10 causes a bulb-like extension at the filopodial tip. This ballooning, spherical model would provide more membrane binding area than the cylindrical model provided here.
